# Joint protein–DNA language modeling enables prediction and engineering of DNA recognition specificity

**DOI:** 10.64898/2026.11.05.763984

**Authors:** Yangfang Xiong, Jiwu Ren, Xiaoxu Wei, Xiaoyu Li, Jiayi Li, Wensheng Wei

## Abstract

Protein–DNA recognition specificity underlies transcriptional regulation, DNA replication, chromatin remodeling and genome editing. Protein–DNA recognition depends on both interacting sequences, yet language models that learn this cross-molecular dependence explicitly during pretraining remain limited. Here, we developed the Protein–DNA Language Models (PDLMs), interaction-aware language models that jointly represent protein and DNA sequences. PDLMs were built by continued pretraining from ESM2 with masked language modeling on paired protein–DNA sequences. A shared encoder places amino acids and nucleotides in the same molecular context, enabling conditional sequence scoring and transferable pair representations. We further introduced a domain-aware masking strategy that increased masking probability within DNA-binding domains to strengthen learning of interaction-relevant regions. Across downstream tasks, PDLMs improved protein–DNA pair classification relative to independent-encoder baselines and supported prediction of mutation-associated binding changes. Zero-shot scoring recovered known changes in DNA-recognition specificity and prioritized variants with higher editing activity. Learned representations also supported cross-study prediction of base editor activity. After fine-tuning on transcription factor binding data, PDLMs predicted DNA motifs directly from protein sequences. Together, PDLMs provide a framework for interaction-aware biomolecular language modeling, with potential applications in protein engineering and genome editing.

## Introduction

Protein–DNA interactions are fundamental to gene regulation and DNA-processing reactions, including replication, recombination, repair and site-specific DNA cleavage^1–5^. Sequence-specific DNA recognition is also central to genome engineering, where DNA-targeting proteins and nucleases determine which genomic sites can be accessed and modified^6–18^. Computational modeling of protein–DNA interactions can provide a quantitative basis for predicting sequence-dependent recognition and guiding the engineering of DNA-targeting proteins. Protein language models provide a scalable framework for learning functional information from sequences. Models including ESM, ProtTrans and ProteinBERT learn transferable representations from large collections of unlabeled protein sequences^19–24^, and their representations and mutation scores have been applied to functional prediction and protein engineering^25–28^. For DNA-binding proteins, these models have been adapted to DNA-related protein functions. ESM-DBP uses domain-adaptive pretraining to improve representations of DNA-binding proteins^29^, while CLAPE^30^ and PDNAPred^31^ use pretrained protein features for DNA-binding site prediction. These approaches strengthen protein-side representations, but do not explicitly model the protein-dependent DNA-recognition specificity.

Recent foundation models have begun to extend sequence modeling across molecular types. LucaOne^32^ and OmniBioTE^33^ learn representations across both sequence types and capture cross-molecular relationships, but do not explicitly model cognate molecular interactions during pretraining. Interaction-conditioned language modeling has developed more directly in protein systems. PLM-interact, MINT and PPLM jointly model interacting proteins and show that partner context can improve interaction-related representations^34–36^. PepMLM further demonstrates target-conditioned sequence modeling for protein–peptide interactions^37^. These studies demonstrate the value of partner-conditioned sequence modeling, but remain within the amino-acid sequence space.

For protein–DNA recognition, the experimental basis for such modeling is increasingly available. Protein-binding microarrays, high-throughput SELEX and related assays have generated large collections of sequence-specific binding measurements^38–42^. Resources such as JASPAR^43^ and CIS-BP^44^ organize experimentally determined binding profiles across diverse transcription-factor families. These data provide a basis for learning protein-dependent DNA recognition. DeepPBS applies bipartite geometric convolutions to protein–DNA complex structures to predict binding specificity^45^. 3M-BSIP uses cross-attention to fuse sequence, structural and surface features for residue-level base-preference prediction^46^. FamilyCode combines a tetrahedral representation of base preferences with family-level PCA and regression to predict mutation-associated specificity changes^47^. Whether partner-conditioned pretraining can be extended across molecular alphabets to learn protein–DNA recognition directly from paired sequences remains unclear. Such pretraining could provide transferable representations of protein–DNA compatibility and recognition specificity for both prediction and engineering.

Here, we developed the Protein–DNA Language Models (PDLMs) by continued pretraining of ESM2 on experimentally characterized protein–DNA binding pairs. Protein and DNA sequences are jointly encoded with modality-specific prediction heads. To emphasize regions involved in recognition, we preferentially masked residues within annotated DNA-binding domains and compared this strategy with uniform masking at two model sizes. We assessed reconstruction on held-out pairs and examined whether the models learned protein-dependent DNA preferences. Conditional scores favored cognate over random DNA and showed a modest association with predicted structural compatibility. We evaluated PDLMs in two modes, zero-shot conditional scoring and transfer of learned representations. In zero-shot analyses, DNA scores recovered a known shift in nickase substrate preference and helped prioritize variants with substrate motif extension. Protein scores also prioritized activity-enhancing substitutions. Frozen joint representations improved pair classification relative to independently encoded features. They also supported prediction of mutation-induced changes in binding free energy. To examine recognition at the nucleotide level, we fine-tuned PDLMs to predict transcription factor binding motifs from protein sequences and recovered a mutation-associated shift in base preference. Overall, these findings support paired pretraining as a framework for modeling partner-dependent recognition and guiding the engineering of protein–DNA interactions.

## Results

### Joint protein–DNA language modeling framework

The PDLM framework was developed to jointly model protein and DNA sequences using paired interaction data (Fig. 1a). We curated 51,329 protein–DNA pairs from BioLiP2^48^, AptaDB^49^, REBASE^50^ and CIS-BP^44^ after sequence filtering and deduplication (Methods). The dataset was divided into training, validation and test sets, with identical protein sequences restricted to a single partition. Each protein–DNA pair was represented as a single input sequence and processed by a shared Transformer initialized from ESM2^20^. Protein and DNA tokens were assigned distinct segment identifiers and learnable segment embeddings to preserve modality information. The protein vocabulary and embeddings were inherited from ESM2^20^. DNA tokens used a separate vocabulary with randomly initialized embeddings. Masked language modeling was applied to both modalities within the same sequence context. Masked amino acids and nucleotides were reconstructed through modality-specific output heads. This design allows protein and DNA tokens to be contextualized jointly while retaining modality-specific prediction spaces. This framework supports zero-shot conditional scoring through the MLM heads and supervised adaptation through frozen representations or task-specific fine-tuning (Fig. 1a).

**Fig. 1.**
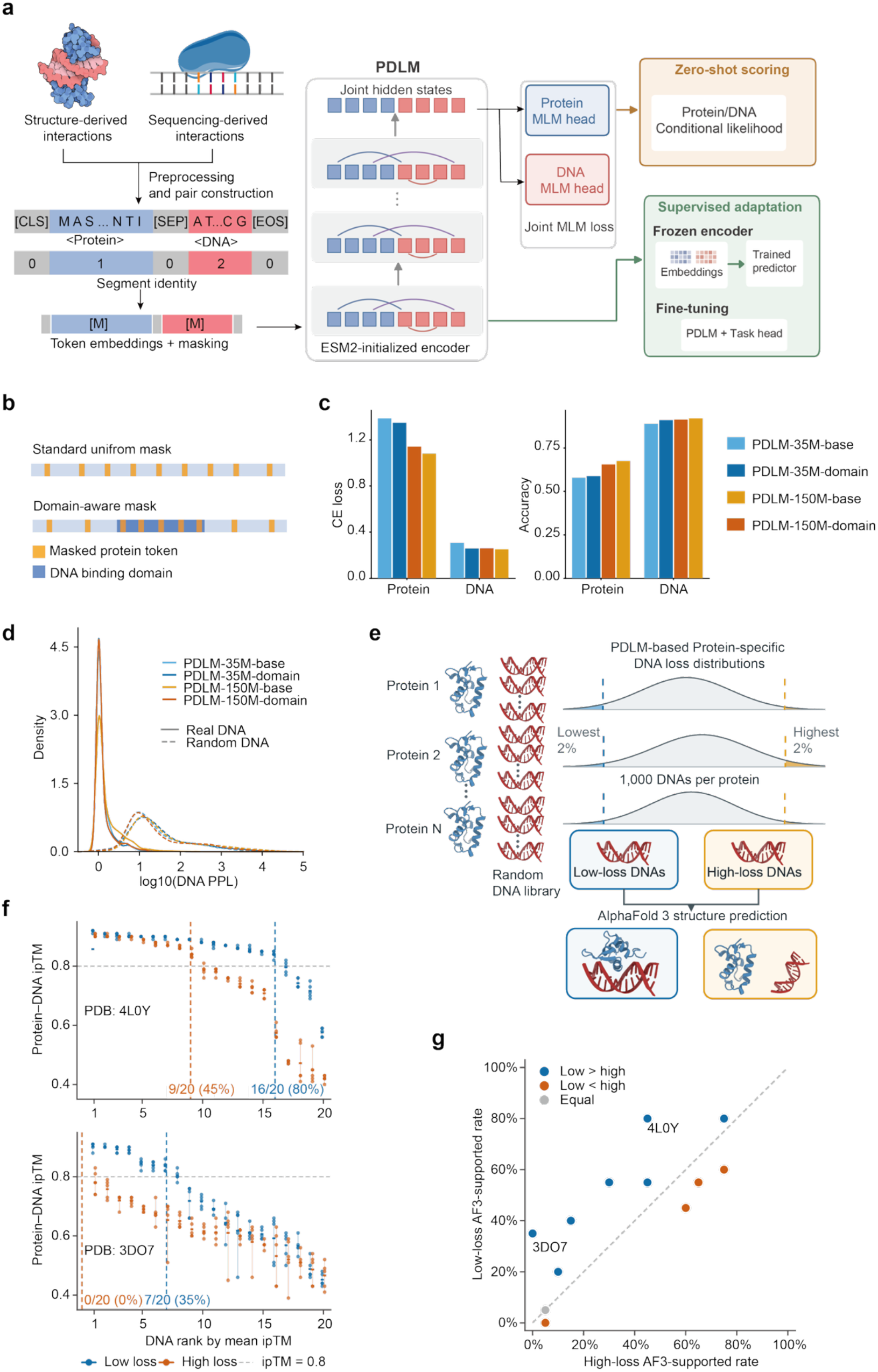
PDLM architecture and evaluation of protein-conditioned DNA modeling. **a**, PDLM architecture. Protein and DNA sequences are jointly processed by a shared Transformer and reconstructed through modality-specific MLM heads. Pretrained models are applied through zero-shot conditional scoring, frozen representations or task-specific fine-tuning. **b**, Uniform and domain-aware protein masking strategies. **c**, Protein and DNA MLM loss and accuracy on the held-out test set. **d**, DNA pseudo-perplexity distributions for cognate and length-matched random sequences. Solid and dashed curves denote cognate and random DNA, respectively. **e**, AF3 evaluation of candidates from the lowest and highest PDLM loss groups. **f**, Protein–DNA ipTM values for two representative proteins. Each point represents an AF3 prediction. Candidates are ranked by mean ipTM. The colored vertical lines and annotations indicate AF3-supported proportions under the criteria in Methods. **g**, Comparison of AF3-supported candidate proportions between the low- and high-loss groups. Each point represents one protein. The diagonal indicates equal proportions in the two groups.

We first optimized the pretraining configuration using an 8-million-parameter ESM2 backbone (Methods). Protein and DNA masking rates were set to 15% and 20%, respectively. We then introduced domain-aware masking to increase the learning signal from protein regions directly involved in DNA recognition. DNA-binding domains were annotated with HMMER using curated Pfam families^51, 52^ (Methods). Residues within annotated DNA-binding domains were masked at a higher probability than residues outside these regions (Fig. 1b).

To distinguish the effects of model scale from those of domain-aware masking, we trained models at two parameter scales under both masking schemes. This yielded four main variants: PDLM-35M-base, PDLM-35M-domain, PDLM-150M-base and PDLM-150M-domain. The base models use uniform protein masking, whereas the domain models use preferential masking of annotated DNA-binding regions. This factorial design allowed us to assess whether gains arose from increased model capacity, domain-focused pretraining or their combination. Training and validation curves of the base models showed stable convergence across model scales.

### Evaluation of protein-conditioned DNA modeling

The four PDLM variants were evaluated by sequence reconstruction on the held-out test set of 2,558 protein–DNA pairs. Protein MLM loss decreased from 1.39 for PDLM-35M-base to 1.08 for PDLM-150M-base, with accuracy increasing from 0.58 to 0.67 (Fig. 1c). DNA reconstruction was less sensitive to model scale, with accuracy ranging from 0.89 to 0.92 across the four variants. Domain-aware masking produced more modest changes in reconstruction performance. At the 35M scale, protein loss decreased from 1.39 to 1.35 and DNA loss from 0.31 to 0.26. Comparable improvements were not observed at the 150M scale. These results indicate that increased model capacity primarily improved protein reconstruction, whereas DNA reconstruction was already comparatively accurate at the smaller scale. Domain-aware masking altered the learning objective without uniformly improving token-level reconstruction, suggesting that its potential benefit should be evaluated through interaction-relevant downstream analyses rather than reconstruction metrics alone.

Token reconstruction, however, does not establish that PDLMs learned protein-dependent DNA preferences. We therefore examined DNA sequence likelihood conditioned on the paired protein (Methods). Cognate DNA sequences showed consistently lower pseudo-perplexity than length-matched random sequences across PDLM variants (Fig. 1d). This indicates that these models capture biologically meaningful sequence patterns beyond random sequence statistics. We next tested whether PDLM-35M-domain sequence scores were associated with an independent measure of protein–DNA structural compatibility using AlphaFold 3 (AF3)^53^. Candidate proteins were randomly sampled from the held-out test set, and 12 proteins whose cognate test-set DNA formed confident AF3-predicted protein–DNA contacts were retained for further analysis. For each protein, we scored 1,000 unique random 15-nt sequences with PDLM-35M-domain and selected the 20 lowest-loss and 20 highest-loss candidates (Fig. 1e). AF3 generated five predictions for each of the resulting 480 protein-DNA duplex complexes. Pairs were classified as AF3-supported using prediction-confidence and contact criteria (Methods). Representative examples showed clear differences between the two score groups. For protein 4L0Y, AF3 supported 80% of low-loss pairs compared with 45% of high-loss pairs. For 3DO7, the corresponding rates were 35% and 0% (Fig. 1f). Across all 12 proteins, seven proteins showed higher support rates in the low-loss group, four showed lower rates and one showed equal rates (Fig. 1g). Thus, low-loss DNA candidates showed a modest tendency toward higher AF3 support, although the association varied across proteins. These results provide independent computational support for an association between PDLM-35M-domain conditional likelihood and predicted structural compatibility, while indicating that the two measures capture related but non-equivalent aspects of protein–DNA recognition.

### Joint representations capture protein–DNA compatibility and mutation effects

To test whether PDLM representations capture pair-specific compatibility, we evaluated protein–DNA binding prediction using experimentally supported pairs and four classes of constructed negatives: mismatched, shuffled and random DNA sequences, and non-DNA-binding proteins (Fig. 2a). Positive pairs retained their original pretraining train, validation and test assignments, and negative pairs were generated within the corresponding partitions, preventing pair-level leakage across splits (Methods). We compared PDLMs with dual-encoder baselines combining ESM2 with either a DNA-6M encoder pretrained separately on our DNA data or DNABERT-2^54^. Both 35M and 150M ESM2 backbones were evaluated. All encoders were frozen, and the same classifier and data splits were used for all models (Methods).

**Fig. 2.**
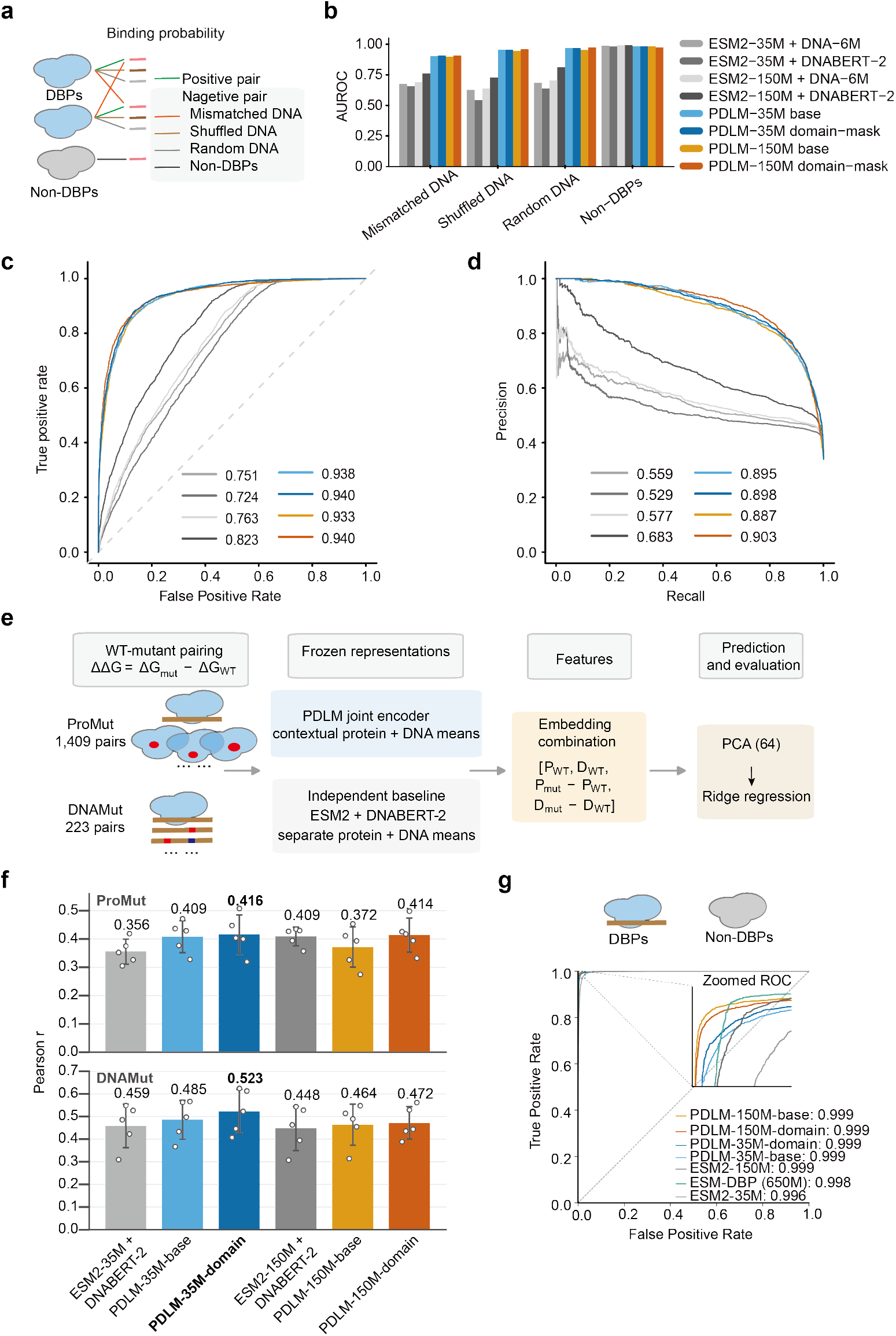
Joint representations for protein-DNA binding and mutation-effect prediction. **a**, Construction of negative protein-DNA pairs using mismatched, shuffled or random DNA sequences and non-DNA-binding protein (non-DBP) controls. **b**, Binding-prediction AUROC by negative-sample type, with each negative subset evaluated against the positive test pairs. **c, d, Overall** receiver operating characteristic (ROC) curves (c) and precision–recall curves (d) across all negative classes. Colors indicate models as in b; values denote AUROC and AUPRC, respectively. **e**, Workflow for ΔΔG prediction using WT representations and WT-to-mutant embedding differences, followed by PCA and ridge regression. **f**, Pearson correlations for protein mutations (top) and DNA mutations (bottom). Bars indicate means across five matched train–test splits, error bars indicate s.d. and points indicate individual splits. **g**, Protein-only DNA-binding protein classification, with an inset showing the low false-positive-rate region. Values denote AUROC.

PDLMs consistently outperformed the dual-encoder baselines on mismatched, shuffled and random DNA negatives, whereas performance on non-DBP negatives was similar across models (Fig. 2b). The same trend was observed in the combined ROC and precision–recall analyses (Fig. 2c, d). PDLM-150M-domain achieved an AUROC of 0.940 and an AUPRC of 0.903, compared with 0.823 and 0.683 for the strongest baseline, ESM2-150M + DNABERT-2. The advantage was therefore most apparent when prediction depended on the compatibility of a specific protein–DNA pair rather than protein-level DNA-binding propensity alone. These results support the benefit of joint protein–DNA representations for pair-specific binding prediction.

We then tested whether these representations also capture quantitative changes in interaction strength. Mutation-induced binding free-energy changes (ΔΔG) were obtained from ProNAB^55^ and analyzed separately for protein mutations and DNA mutations (Fig. 2e). The resulting dataset contained 1,409 protein-mutant pairs and 223 DNA-mutant pairs. For each example, the feature vector combined the wild-type protein and DNA representations with the corresponding mutant-minus-wild-type representation difference. The same feature construction was applied to independently encoded ESM2 and DNABERT-2 representations. Principal-component analysis was followed by ridge regression. The ridge penalty was selected by five-fold cross-validation within each training set, and performance was summarized across five matched repeated random 8:2 pair-level train-test splits (Methods).

PDLM-35M-domain achieved the highest mean Pearson correlation for both mutation classes (Fig. 2f). For protein mutations, its performance was similar to that of the strongest independent baseline (0.416 versus 0.409). A larger difference was observed for DNA mutations, for which PDLM-35M-domain reached 0.523 compared with 0.459 for the strongest baseline. The advantage of joint representations was therefore more pronounced for DNA sequence perturbations in this benchmark, consistent with the explicit contextualization of DNA by its protein partner.

As a complementary test, we asked whether joint pretraining preserved protein-intrinsic information associated with DNA binding. In protein-only DNA-binding protein classification, PDLMs, ESM-DBP^29^ and ESM2 all achieved near-ceiling performance, with AUROC values of 0.996–0.999 (Fig. 2g). The saturation of this benchmark limits meaningful ranking among the models. Instead, the result shows that joint protein–DNA pretraining preserves protein-level features sufficient for DNA-binding protein classification. Together with the pairwise binding and mutation-effect analyses, these results indicate that PDLMs retain protein-intrinsic information while adding representations that are sensitive to the identity and sequence of the DNA partner.

### Zero-shot protein-conditioned DNA scoring captures MutH substrate preference and guides variant screening

The pretraining corpus included diverse nuclease–DNA recognition pairs, providing a basis to test whether PDLMs could transfer this information to zero-shot specificity prediction. We focused on MutH, the sequence-specific nickase used in mitochondrial base editors (mitoBEs)^11^, where its recognition preference constrains the accessible editing sites. Wild-type MutH recognizes 5′-GATC-3′^56^, whereas the engineered E91A/F94A variant broadens editing to GATN contexts, increasing the number of candidate sites from 23 to 485 ^11^. We therefore asked whether PDLMs could recover this established specificity shift without task-specific training. For each GATN sequence, we calculated the change in protein-conditioned DNA pseudo-log-likelihood between E91A/F94A and wild-type MutH (Fig. 3a, b). Positive values indicate increased model preference for the same DNA sequence in the mutant background. Among the four PDLM variants, PDLM-35M-domain assigned positive changes to all three alternative substrates, GATA, GATG and GATT, while the GATC score remained nearly unchanged (Fig. 3c). Thus, zero-shot conditional scoring recovered the direction of the known substrate expansion and provided a basis for further variant screening.

**Fig. 3.**
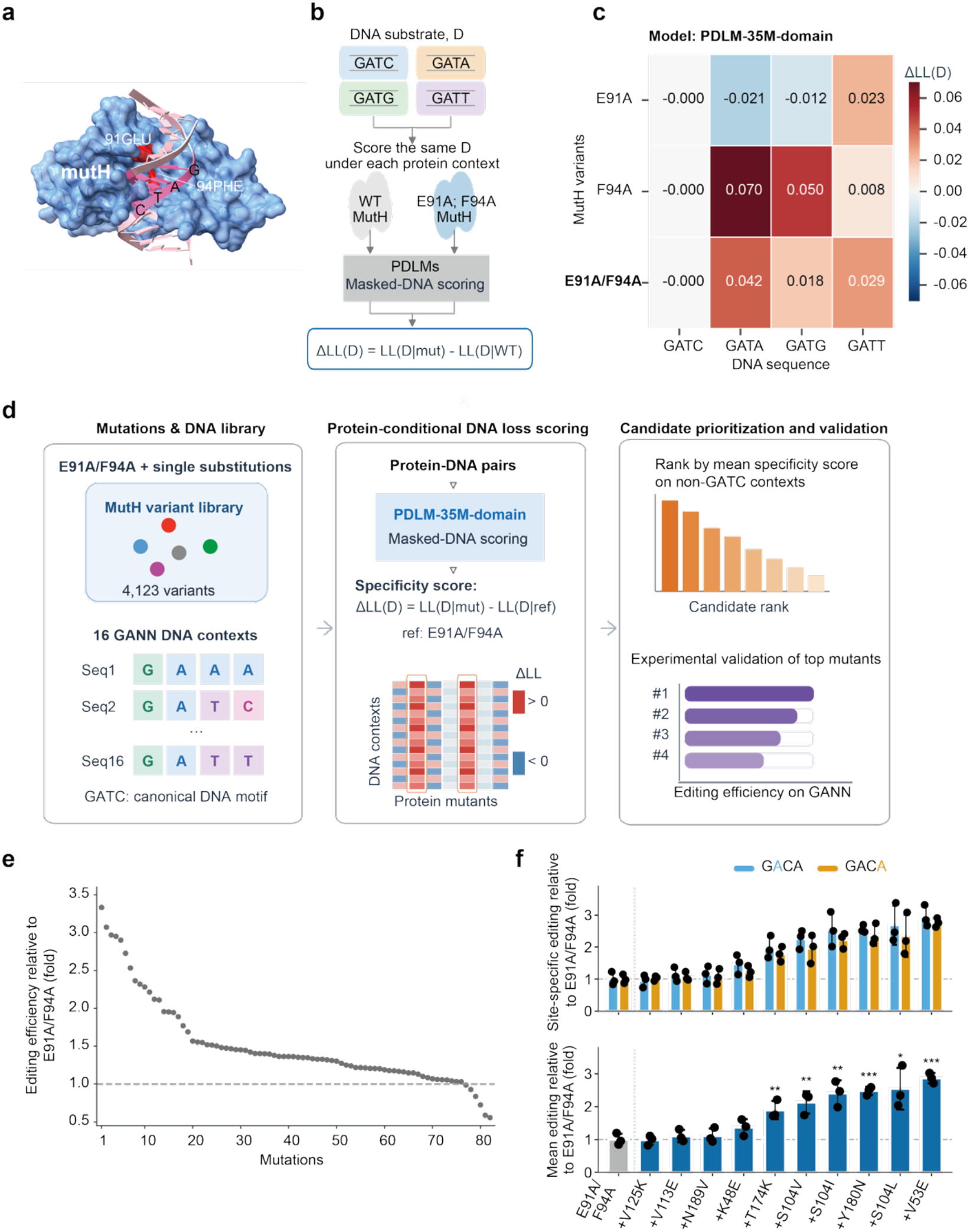
PDLM recovers MutH specificity changes and guides substrate expansion. **a**, Structure of MutH bound to DNA, highlighting residues E91 and F94. **b**, Protein-conditioned DNA scoring of the known GATN specificity change, using wild-type MutH as the reference. **c**, Changes in DNA pseudo-log-likelihood for E91A, F94A and E91A/F94A across the four GATN substrates using PDLM-35M-domain. Positive values indicate increased model preference in the mutant protein context. **d**, Workflow for model-guided MutH variant screening. A library of 4,123 single substitutions in the E91A/F94A background was scored across all 16 GANN contexts. Candidates with positive score changes across all 15 non-GATC contexts were prioritized by their mean score change for experimental evaluation. **e**, Editing efficiencies of the 82 prioritized variants at GACA, shown relative to the E91A/F94A background. **f**, Follow-up validation of ten selected MutH variants at GACA. Blue and orange indicate editing at the first and second adenines, respectively; the lower panel shows their arithmetic mean. Bars show mean ± s.d. “+” denotes an additional substitution in the E91A/F94A background. Asterisks indicate significance by Student’s t-tests versus parental E91A/F94A (*P < 0.05, **P < 0.01, ***P < 0.001).

Further expansion of MutH recognition could increase the sequence space accessible to mitoBEs. The mitoBE system currently lacks a practical high-throughput screening framework for large-scale variant optimization. Because individual MutH variants must be constructed separately and evaluated using site-specific cellular editing assays, systematic mutational screening remains experimentally intensive and difficult to scale. Computational prioritization therefore provides a practical approach to narrow the experimental search space and focus validation on the most promising candidates. We used PDLM-35M-domain to score 4,123 additional single substitutions in the E91A/F94A background across all 16 GANN contexts (Fig. 3d). E91A/F94A served as the reference, so each score quantified the predicted effect of an additional substitution beyond the established double mutant. Candidates were required to show positive score changes across all 15 non-GATC contexts and were ranked by their mean score change. This procedure prioritized 82 variants for experimental evaluation.

We next tested whether PDLM-guided variant prioritization could improve editing at GACA, a non-GATN sequence context, using the mitoBE system. All 82 variants were experimentally evaluated in two screening rounds, comprising 30 and 52 variants, respectively. The ten highest-performing variants were subsequently evaluated in three independent biological replicates. Six variants showed significantly increased mean editing relative to E91A/F94A (Fig. 3f). Collectively, these results indicate that PDLM-35M-domain captures protein-dependent DNA-recognition specificity and can translate this information into zero-shot virtual screening of MutH variants.

### Protein-side zero-shot scoring prioritizes activity-enhancing MutH substitutions

After using protein-conditioned DNA scores to model MutH substrate specificity, we next examined the complementary protein-side application of PDLMs for activity optimization. With GATA fixed as the substrate, we tested whether protein-only zero-shot scores could identify substitutions that improve editing activity in the E91A/F94A background. We evaluated 179 single amino-acid substitutions with measured GATA editing activities (Methods). Each substitution was scored from masked amino-acid probabilities without DNA input or task-specific fitting (Fig. 4a).

**Fig. 4.**
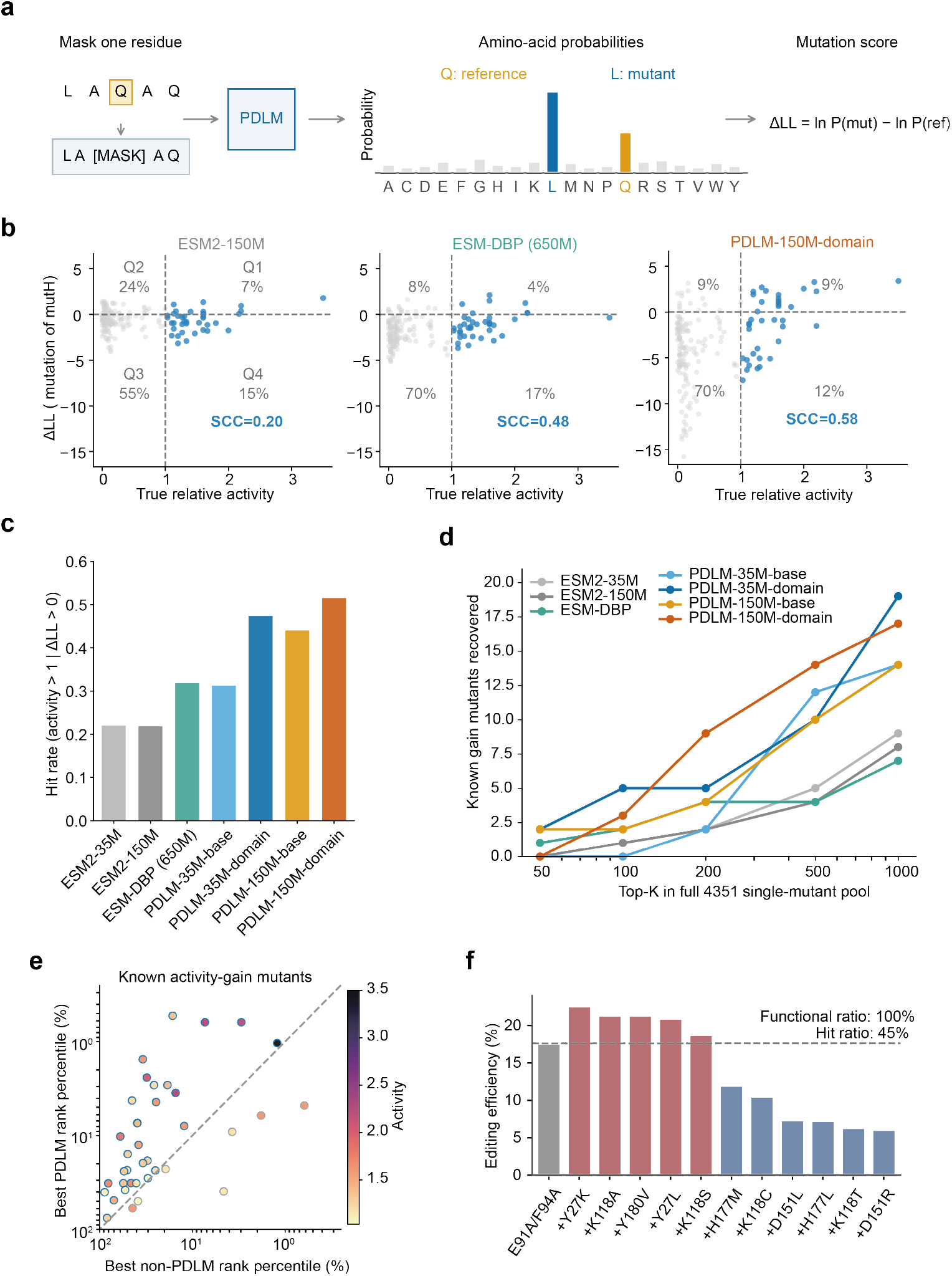
PDLM scores recover and prioritize MutH activity-enhancing substitutions. **a**, Protein-only masked amino-acid scoring of MutH substitutions. **b**, Protein ΔLL plotted against relative GATA editing activity. Blue points denote activity-enhancing variants. Dashed lines mark reference activity and zero score. Quadrant percentages use all tested variants, whereas SCC is calculated within the enhancing subset. **c**, Fraction of activity-enhancing variants among tested substitutions with positive scores. **d**, Recovery of known enhancing variants within top-ranked subsets of the full single-substitution library. **e**, Best rank percentiles within the PDLM and baseline model groups for enhancing variants. **f**, Experimental validation of 11 substitutions selected by PDLM consensus ranking. Candidates were required to rank within the top 50 in at least three of the four PDLM variants.

Among activity-enhancing substitutions, the SCC between model scores and measured GATA editing activity was 0.58 for PDLM-150M-domain, compared with 0.48 for ESM-DBP and 0.20 for ESM2-150M (Fig. 4b). The domain-aware PDLM variants also showed higher fractions of activity-enhancing substitutions among positively scored variants than the baselines (Fig. 4c). Across the full single-substitution space, all four PDLM variants recovered more known activity-enhancing substitutions than the three baselines at both the top-500 and top-1,000 cutoffs (Fig. 4d). The best PDLM rank was also higher than the best baseline rank for most substitutions with increased activity (Fig. 4e).

We then used consensus ranking for experimental candidate selection. Eleven substitutions ranked within the top 50 in at least three of the four PDLM variants were tested. All 11 retained editing activity, and five showed higher GATA editing than the parental E91A/F94A variant (Fig. 4f), corresponding to functional and activity-improvement ratios of 100% and 45%, respectively. The fraction of activity-improving variants was comparable to that observed among positively scored substitutions in the initial dataset. These results show that protein-side PDLM scoring can enrich functional and activity-improving mutations and support efficient candidate prioritization across a large protein sequence space.

### PDLMs support cross-dataset TadA activity prediction and zero-shot off-target scoring

To test whether PDLM representations transfer to a distinct DNA-editing system, we next examined TadA base editors. Two independently generated datasets provided a stringent cross-study setting because they contained no identical TadA variants or DNA target sequences. We trained support vector regression models on the Xiao et al. dataset using protein-conditioned DNA representations from the five-nucleotide window centered on each edited adenine (Fig. 5a)^57^. In fivefold internal cross-validation, all four PDLM variants achieved modestly higher mean SCCs (0.87–0.90) than the baselines that encoded TadA and DNA separately with ESM2 and DNABERT-2^54^ (Fig. 5b). PDLM-150M-base achieved an SCC of 0.90 across the pooled out-of-fold predictions (Fig. 5c). We then evaluated the trained models on the smaller, independent Chen et al. dataset^58^, where all four PDLM variants retained positive correlations (SCCs of 0.56–0.62), whereas both baselines yielded negative correlations (Fig. 5b). These results indicate that PDLM representations capture transferable features of TadA editing activity across protein variants and DNA sequence contexts.

**Fig. 5.**
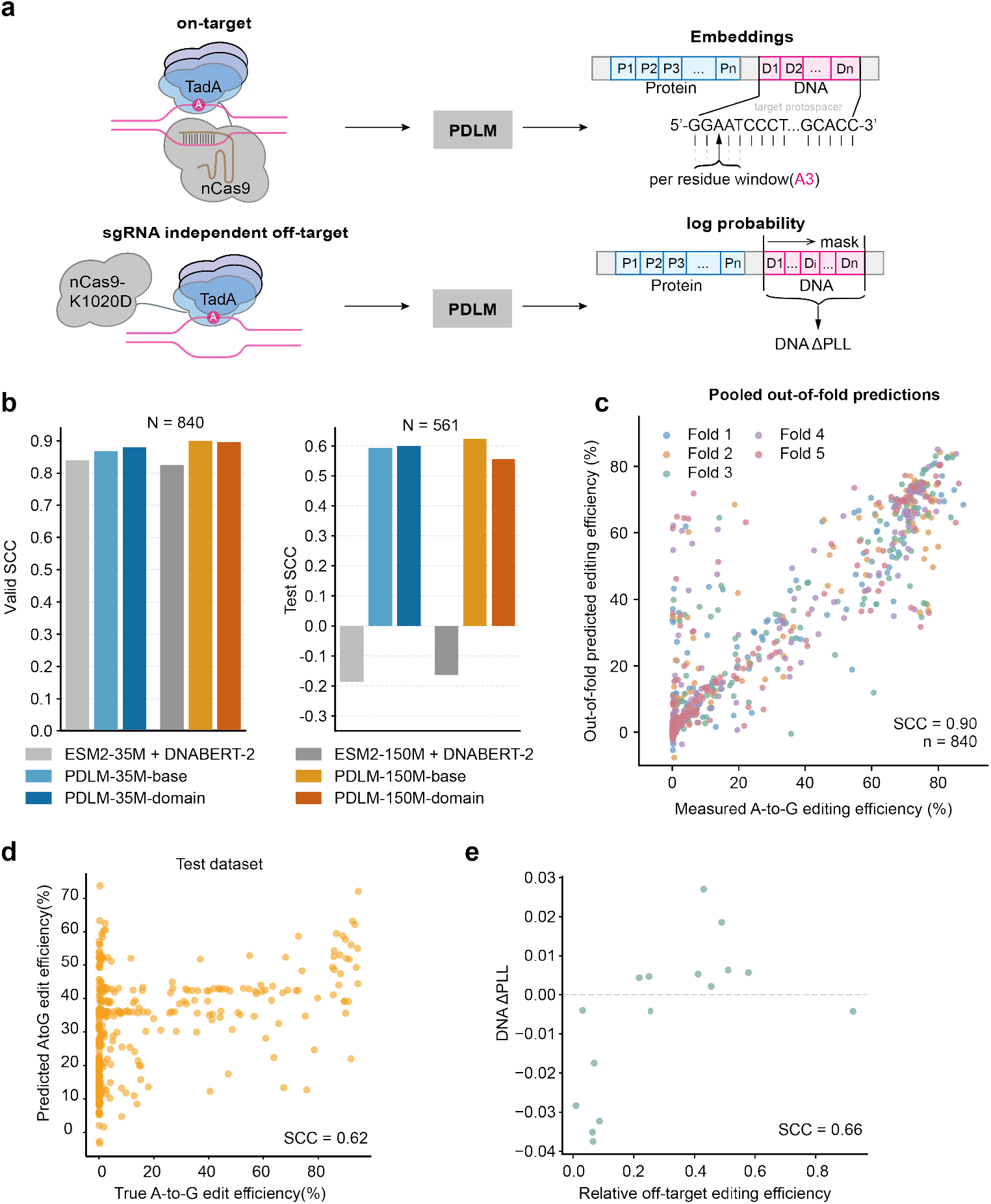
PDLMs support on-target activity prediction and off-target scoring of TadA variants. **a**, Schematic of supervised on-target activity prediction from local protein-conditioned DNA representations and zero-shot scoring of sgRNA-independent DNA off-target activity. **b**, TadA on-target activity prediction using four PDLM variants. Left, mean Spearman correlation coefficient (SCC) across internal cross-validation folds in the Xiao et al. dataset. Right, SCC on the independent Chen et al. dataset. The independently encoded ESM2 + DNABERT-2 baselines were trained and evaluated using the same procedure. **c,** Predicted versus measured A-to-G editing efficiencies for PDLM-150M-base, pooling out-of-fold predictions from all five internal cross-validation folds (n = 840). Each point represents a single adenine in a given TadA variant–DNA target pair and is colored by its held-out fold. **d,** Predicted versus measured A-to-G editing efficiencies for PDLM-150M-base on the independent Chen et al. dataset^58^. **e**, Change in protein-conditioned DNA pseudo-log-likelihood plotted against relative sgRNA-independent off-target editing efficiency. Scores were calculated with PDLM-150M-domain relative to TadA8e. Each point represents one TadA variant.

We next asked whether protein-conditioned DNA scoring could capture TadA-dependent off-target activity without task-specific fitting. We evaluated 16 non-reference TadA variants on the same DNA substrate in an orthogonal R-loop assay^59^, providing a matched sequence context for comparison. PDLM-150M-domain scores measured the change in mean DNA pseudo-log-likelihood relative to TadA8e on that substrate. These scores were positively associated with relative off-target editing (SCC = 0.66; Fig. 5e). Thus, protein-conditioned DNA likelihood provides a complementary zero-shot measure of TadA-dependent editing behavior beyond supervised on-target prediction. In contrast, protein-only mutation log-odds showed weak and inconsistent correlations across ESM2 and PDLM variants, whereas protein-conditioned DNA scores showed stronger associations in the best-performing models. These results suggest that partner-conditioned DNA scoring captures information relevant to TadA-dependent off-target activity beyond protein mutation likelihood alone.

Overall, the TadA analyses show that paired protein–DNA pretraining can support both cross-dataset activity prediction and zero-shot scoring of TadA-dependent off-target effects.

### Fine-tuning adapts PDLMs for transcription factor motif prediction

Transcription factor motif prediction provided a complementary setting to assess whether PDLM representations could be adapted to nucleotide-level binding specificity. Protein sequences and experimentally determined position weight matrices (PWMs) were obtained from CIS-BP^44^. Protein–PWM pairs inherited the protein-sequence-level train, validation and test assignments used for pretraining, yielding 7,129 training, 396 validation and 396 test pairs (Methods). During fine-tuning, all DNA positions were masked, and a position-wise prediction head converted the corresponding DNA representations into nucleotide probabilities (Fig. 6a). Using PDLM-35M-domain, we first compared different extents of encoder updating. Fine-tuning the final one-third of the encoder increased the median PWM Pearson correlation coefficient (PCC) from 0.40 to 0.64 relative to a frozen encoder and reduced the corresponding prediction error (Fig. 6b). This partial fine-tuning scheme was therefore used for subsequent comparisons.

**Fig. 6.**
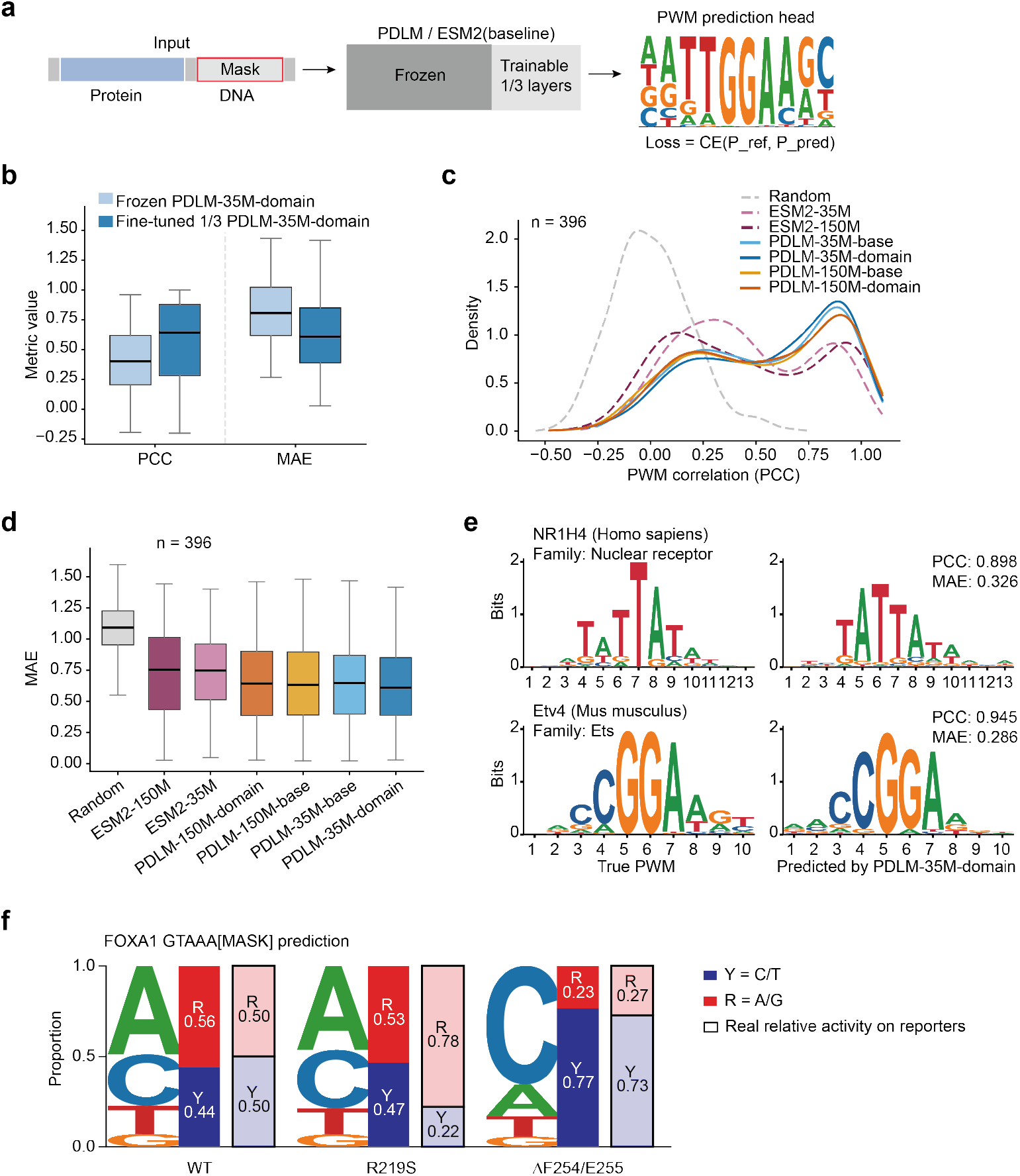
Fine-tuned PDLMs predict transcription factor DNA-binding motifs. **a**, PWM prediction from protein sequences and fully masked DNA positions. The prediction head and final one-third of the encoder layers are updated during fine-tuning. **b**, PWM correlation and prediction error for PDLM-35M-domain with a frozen or partially fine-tuned encoder. **c, d,** Distributions of per-motif Pearson correlation coefficients (c) and PWM prediction errors (d) for PDLM variants, ESM2 baselines and a random PWM baseline. **e**, Experimental and predicted sequence logos for NR1H4 and Etv4. **f**, Predicted preferences for purines (R, A/G) and pyrimidines (Y, C/T) at the sixth motif position for wild-type FOXA1, R219S and the F254–E255 deletion, compared with normalized reporter activities.

Under the same training procedure, PDLM-35M-domain achieved the highest median PWM correlation and the lowest median error among the evaluated PDLM and ESM2 models (Fig. 6c, d). Increasing model size to 150M parameters did not improve performance with the available training data. Representative predictions for NR1H4 and Etv4 reproduced the major nucleotide preferences of the experimental motifs (Fig. 6e). These results show that PDLM representations can be adapted to infer protein-dependent DNA-binding motifs directly from protein sequence.

We further examined mutation-associated changes in transcription factor specificity using published FOXA1 reporter measurements^60^. R219S and the F254–E255 deletion shift nucleotide preference at the variable position immediately following GTAAA. We therefore predicted nucleotide probabilities at this position from the wild-type and mutant protein sequences. The predicted shift toward pyrimidines for the F254–E255 deletion closely matched the experimental trend. The model did not recover the experimentally observed shift toward purines for R219S (Fig. 6f). Overall, fine-tuning extended PDLMs from general protein–DNA representations to direct prediction of transcription factor motifs and mutation-associated changes in DNA preference.

## Discussion

In this study, we developed PDLMs to incorporate protein–DNA interaction information directly into language-model pretraining. The central advance lies in using molecular partnerships as the learning context, allowing sequence variation to be evaluated in relation to an interacting partner. We can use each pretrained model in two complementary ways. Conditional likelihoods enable zero-shot exploration of protein or DNA sequence space, whereas learned representations can be adapted to supervised prediction when labeled data are available. Across the analyses presented here, these two modes captured different aspects of protein–DNA recognition, including partner compatibility, sequence preference and mutation-associated functional variation. The MutH application further shows that conditional scoring can be used to reduce a large mutational search space and enrich variants with altered recognition or improved editing activity. These results support paired pretraining as a useful basis for both predictive modeling and sequence-guided protein engineering.

The four PDLM variants show that increased model size and domain-aware masking provide complementary rather than uniform benefits. PDLM-35M-domain performed strongly in several recognition-specificity tasks, including MutH substrate scoring, mutation-associated binding prediction and transcription-factor motif prediction. Larger models showed clearer advantages in some protein-side and interaction tasks, with PDLM-150M-domain performing strongly in protein mutation scoring, pairwise binding prediction and TadA off-target analysis. PDLM-150M-base also showed the strongest cross-dataset TadA activity prediction. These differences indicate that scaling alone does not determine downstream performance at the present data scale. Domain-aware masking can increase sensitivity to recognition-related sequence features, but its benefit depends on the task and model size. Maintaining models with different capacities and masking objectives may therefore be more useful than selecting a single universal configuration.

PDLMs are currently limited by the scale and diversity of available paired protein–DNA data. Experimentally characterized protein–DNA pairs are far fewer than protein-only sequences and are unevenly distributed across protein families and assay types. This uneven coverage may restrict transfer to underrepresented recognition systems, and generalization to distant homologues and unseen protein families remains to be established. Quantitative datasets that jointly sample variation on both sides of the interaction would be particularly valuable. Protein-by-DNA specificity matrices, systematic protein mutagenesis and matched affinity measurements could directly capture how protein sequence changes reshape DNA preference. Such quantitative paired datasets remain limited and will be important for future development of conditional scoring and specificity-oriented protein design.

PDLMs provide a starting point for incorporating molecular interaction information directly into language-model pretraining. By making the interacting partner part of the learning context, paired pretraining connects sequence representation with partner-dependent function and protein engineering. Future models trained on broader quantitative interaction datasets and complementary structural information could extend this approach to more diverse recognition systems. More broadly, learning from molecular partnerships offers a route toward biomolecular models that predict how sequence variation reshapes interactions and guide their engineering.

## Methods

### Protein–DNA pretraining data

Protein–DNA pairs from BioLiP2^48^, REBASE^50^, CIS-BP^44^ and AptaDB^49^ were filtered and deduplicated. REBASE proteins were clustered with MMseqs2^61^ at sequence identity and coverage thresholds of 0.9. Retained pairs contained proteins of 50–1,000 residues and DNA of 3–50 nucleotides. The 51,329 pairs were split into 46,224 training, 2,547 validation and 2,558 test pairs, keeping identical proteins within one partition (seed 42).

### PDLM architecture and pretraining

A shared ESM2-initialized Transformer encoded “[CLS] protein [SEP] DNA [EOS]” using separate vocabularies and learned segment embeddings. Protein embeddings were inherited from ESM2; DNA embeddings and separate linear MLM heads were newly initialized. Inputs were capped at 800 tokens by trimming proteins while retaining DNA.

Uniform masking selected 15% of protein and 20% of DNA tokens, with equally weighted mean cross-entropy losses. Domain-aware masking used HMMER/Pfam^51, 52^ annotations: protein masking probabilities were 0.30 inside DNA-binding regions, 0.10 outside and 0.15 for unannotated proteins. DNA masking remained at 20%. Reverse-complement augmentation was applied to DNA with probability 0.30.

The 35M and 150M models were trained at a learning rate of 4 × 10^−4^ for up to 40 epochs with validation-based early stopping. Held-out reconstruction loss and accuracy used uniform masking. Seed 42 was used unless specified.

### DNA scoring and AF3 evaluation

Masking each DNA base with the remaining sequence and protein visible gave the mean DNA pseudo-log-likelihood and reference-relative change:

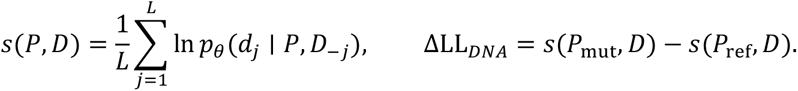

*D*_−j_ masks position *j*; DNA loss was −*s*(*P*, *D*) and pseudo-perplexity was exp[−*s*(*P*, *D*)]. Cognate and length-matched random DNA were compared under the same protein context.

Twelve randomly sampled held-out proteins passed cognate-DNA AF3 contact screening^53^. PDLM-35M-domain scored 1,000 unique random 15-nt sequences per protein. The 20 lowest- and 20 highest-loss sequences, paired with their reverse complements, yielded 480 duplex complexes for AlphaFold Server prediction^53^ (five models each).

Protein–DNA ipTM used the larger protein–strand value; PAE used the smaller. Supported models required ipTM > 0.8, PAE ≤ 10 Å, ≥20 protein–DNA atom pairs within 5 Å and ≥3 DNA residues contacting the protein within 5 Å. Pairs required support from ≥3 of five models. Pooled pair counts and paired protein-level support rates were compared using two-sided Fisher’s exact and Wilcoxon signed-rank tests, respectively.

### Protein–DNA binding prediction

Pretraining partitions were retained, with approximately two within-partition negatives per positive: mismatched DNA (50%), shuffled DNA (12.5%), random DNA (12.5%) and non-DNA-binding proteins paired with DNA (25%). DNA controls matched length and, where applicable, composition or GC content; known positives were excluded. Training, validation and test sets contained 136,675, 7,502 and 7,539 pairs, with no identical proteins or pairs shared across partitions.

Frozen PDLM features concatenated CLS and mean protein and DNA embeddings. Baselines independently encoded partners using ESM2-35M or ESM2-150M with DNA-6M or DNABERT-2^54^. All models shared the classifier and splits; only classifiers were trained. AUROC and AUPRC were evaluated on the complete test set and for each negative class against all 2,558 positive test pairs.

### Prediction of mutation-associated binding free-energy changes

ProNAB^55^ WT–mutant pairs targeted ΔΔ*G* = Δ*G*_mut_ − Δ*G*_WT_ . Annotation errors, invalid measurements and inputs >800 tokens were excluded; WT free energies had to match *RT*ln*K_d_* within 0.2 kcal mol⁻¹. Identical sequence pairs sharing an experimental-condition identifier were aggregated by median ΔΔG, yielding 1,409 protein-mutant and 223 DNA-mutant pairs, analyzed separately.

Mean protein and DNA embeddings excluded special/padding tokens. PDLM jointly encoded partners; the baseline used independent ESM2 and DNABERT-2 encoders. Features combined WT embeddings with mutant-minus-WT differences:

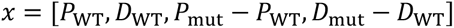

Randomized PCA reduced features to 64 dimensions without prior standardization. Ridge regularization was selected from 10⁻⁴ to 10⁶ by five-fold inner cross-validation, with PCA refitted within folds. Five matched 8:2 pair-level splits (seeds 42–46) retained assignments after single-condition filtering. Outer tests reported Pearson correlation, RMSE and MAE.

### DNA-binding protein classification

Linear classifiers used protein-only PDLM, ESM2 and ESM-DBP embeddings for 26,672 BioLiP2^48^ positives and 26,672 UniRef50^62^ negatives lacking nucleic-acid-binding annotations and DNA-binding-related domains. Stratified five-fold cross-validation provided out-of-fold ROC curves.

### MutH specificity screening and GACA validation

Wild-type MutH, E91A, F94A and E91A/F94A were scored against GATC, GATA, GATG and GATT. Reference proteins were wild-type MutH for GATN validation and E91A/F94A for additional-substitution screening.

PDLM-35M-domain scored 4,123 additional substitutions against 16 GANN sequences, excluding positions 91 and 94 and the first and last five residues. Positive Δ*LL*_DNA_ across all 15 non-GATC sequences selected 82 variants, ranked by mean score change. Residue pLDDT and DNA distance prioritized 30 for first-round testing; 52 were tested subsequently.

GACA editing of all 82 variants was normalized to within-round E91A/F94A controls. Ten selected variants underwent validation in three independent biological replicates, reported as mean ± sample s.d. Comparisons with E91A/F94A used unpaired, two-sided Student’s t-tests.

### MutH activity scoring and candidate validation

The 179 single substitutions in E91A/F94A were characterized for GATA activity before PDLM development in a MutH campaign including EVOLVEpro-guided prioritization^25^. Four PDLM variants, ESM2-35M, ESM2-150M and ESM-DBP were evaluated without task-specific fitting or DNA input. Masking each reference residue gave:

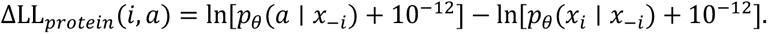

Both probabilities came from the same masked input after vocabulary-wide softmax. SCC was calculated among the 38 substitutions with increased activity. Hit rate was the fraction with activity > 1 among tested variants with positive scores. All 4,351 substitutions were ranked to assess recovery of these 38 substitutions at top-K cutoffs. The best rank percentiles from four PDLM variants were compared with those from three baselines. Eleven candidates in the top 50 of ≥3 PDLM variants were tested, normalizing editing to the assay reference.

### TadA on-target prediction and off-target scoring

Predictors used Xiao et al.^57^ data (n = 840) for training and Chen et al.^58^ data (n = 561) for external tests. Duplicate measurements were averaged. Support vector regression used frozen PDLM DNA embeddings from joint protein–DNA inputs, averaged over five nucleotides centered on the edited adenine. Five-fold internal cross-validation grouped identical full-protein and normalized-DNA pairs. Independent-encoder baselines used ESM2-35M or ESM2-150M with DNABERT-2 and shared the DNA window, regression procedure and splits. SCC assessed performance.

Sixteen non-reference TadA variants from an orthogonal R-loop assay^59^ were scored on a fixed DNA substrate. Editing was normalized to TadA8e. DNA scores were the change in mean protein-conditioned DNA pseudo-log-likelihood relative to TadA8e, calculated by masking each nucleotide in turn. Positive changes indicated higher likelihood under the mutant protein context. SCC assessed agreement with relative off-target editing. For comparison, protein-only mutation scores relative to TadA8e were calculated as described above and correlated with relative off-target editing across the same variants.

### Transcription factor motif prediction

CIS-BP^44^ protein–PWM pairs retained pretraining partitions (7,129 training, 396 validation, 396 test). Proteins were paired with fully masked DNA of target motif length. We trained a linear head with a softmax output using position-wise cross-entropy. We compared a fully frozen encoder with one in which the final third of the layers was fine-tuned. Head and backbone learning rates were 2 × 10⁻⁴ and 1 × 10⁻⁵, respectively, with batch size 128, weight decay 0.01 and warm-up ratio 0.1. Training ran for up to 20 epochs with an early-stopping patience of three epochs. Checkpoints were selected by the validation score (correlation + 0.25 accuracy − 0.10 KL − 0.05 loss).

PCC compared flattened predicted and target PWMs; the higher-PCC target orientation was also used for error:

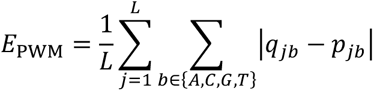

MAE summed absolute errors over four bases, then averaged over positions. Random PWMs were sampled from a Dirichlet(1,1,1,1) distribution using seed 42.

FOXA1 wild-type, R219S and F254–E255 deletion variants were paired with GTAAA[MASK]. Predicted purine and pyrimidine probabilities were compared with GTAAAR and GTAAAY reporter activities after normalization by their sum^60^.

### MutH specificity validation and editing analysis

HEK293T cells transfected with MutH and TadA plasmids were collected after 72 h for genomic DNA extraction and targeted amplicon sequencing. Illumina HiSeq X Ten reads were aligned with BWA^63^ and sorted with SAMtools^64^. REDItools^65^ detected targeted base conversions by Fisher’s exact test (P < 0.05).

## Data availability

Source data underlying the figures and the processed evaluation datasets will be available at https://github.com/xiongyf520/PDLM. Publicly available data used in this study were obtained from REBASE (https://rebase.neb.com/rebase/rebase.html), BioLiP2 (https://zhanggroup.org/BioLiP/), AptaDB (https://lmmd.ecust.edu.cn/aptadb/), CIS-BP (https://cisbp.ccbr.utoronto.ca/), ProNAB (https://web.iitm.ac.in/bioinfo2/pronab/) and UniRef50 (https://www.uniprot.org/uniref). Published TadA editing and FOXA1 reporter data were obtained from the studies cited in Methods. Dataset selection and preprocessing procedures are described in Methods.

## Code availability

Code and configuration files for PDLM pretraining, zero-shot scoring and representation extraction are available on GitHub at https://github.com/xiongyf520/PDLM. Model weights are available on Hugging Face at https://huggingface.co/xiongyf/PDLM.

## Acknowledgements

W.W. received funding from the National Natural Science Foundation of China (32530017).Computational resources were provided by the High-Performance Computing Platform of Peking University. We thank Dr. Z.Y. for valuable discussions during the early stage of this project, particularly on DNA-binding specificity, and for insightful guidance on the optimization of MutH.

## Author contributions

The work presented in this paper was a collaborative effort of all authors. W.W. supervised the project. Y.X. and J.R. conceived the study and designed the model architecture. Y.X. conducted model training and fine-tuning. Y.X., J.R. and X.L. performed downstream evaluations and application studies. X.W. conducted the experimental validation of MutH with assistance from J.L. X.W. and J.R. performed editing efficiency analysis. Y.X. prepared the initial manuscript and figures. J.R., X.L., and X.W. contributed to data analysis, figure preparation, and manuscript writing. W.W. reviewed and revised the manuscript. All authors read and approved the final manuscript.

